# Boron-induced microbial filtering delays early rhizosphere establishment in agricultural soil

**DOI:** 10.64898/2026.09.08.750147

**Authors:** Subhajit Sen, Nibendu Mondal, Chandana Basak, Wriddhiman Ghosh, Ranadhir Chakraborty

**Author notes:** Corresponding author: Prof. Ranadhir Chakraborty, Postal address: OMICS Laboratory, Department of Biotechnology, University of North Bengal, Siliguri- 734013, West Bengal, India. CSIR-Centre for Cellular and Molecular Biology (CCMB), Hyderabad, Telangana, India. Department of Microbiology & Cell Science, Fort Lauderdale Research and Education Center, University of Florida, Davie, Florida 33314, USA. Microbial Research Centre, BRIC-Translational Health Science and Technology Institute (THSTI), Faridabad, India.

## Abstract

Excessive application of boron-based fertilizers in agriculture has led to increased boron accumulation in soil, adversely affecting microbial communities and plant species, ultimately contributing to boron-induced desertification. This study presents the first metagenome-based analysis of boron-contaminated agricultural soil in Northern West Bengal, India, providing a comprehensive profile of the soil microbiome under boron stress. Boron-amended and unamended agricultural soil samples were analyzed using metagenomics sequencing approach over 365 days under natural environmental conditions. The results revealed a significant decline in bacterial diversity in boron-amended soil, with Firmicutes emerging as the dominant phylum. Notably, all boron-tolerant bacterial strains belonged to *Firmicutes*, suggesting that high boron levels impose selective pressure favouring Gram-positive bacteria. The enrichment of metal-resistant and oligotrophic bacterial taxa in boron-amended soil underscores microbial adaptation to boron stress. In addition to microbial community dynamics, we investigated plant succession in boron-enriched soil, offering novel insights into the interplay between above- and below-ground ecosystems. Our findings highlight two key adaptive traits, high boron tolerance and oligotrophic survival strategies, that enable bacterial communities to persist in boron-rich environments. These results enhance our understanding of microbial resilience in chemically stressed soils and have implications for soil health and sustainable agriculture.

## 1. Introduction

Boron, a group 13 metalloid with atomic number 5 and atomic mass 10.811 u, is widely distributed in nature, occurring in sedimentary rocks (∼85 ppm), soils (∼0.2–30 ppm), coal (∼5–400 ppm), and seawater (∼4.6 ppm) (Harder, 1970; Swain, 1994; Swain and Goodarzi, 1995; Brdar-Jokanović, 2020). In soils, boron exists primarily as boric acid [B(OH)] (∼96%) and to a lesser extent as the borate anion [B(OH)], playing a crucial role in various metabolic and physiological processes in plants, animals, and microorganisms (Lovatt and Dugger, 1984; Lou et al., 2001; Bolaños et al., 2004). While essential for plant growth, boron has a narrow range between deficiency and toxicity, making its management in agricultural soils particularly challenging (Paull et al., 1991). Boron toxicity arises when soil concentrations exceed 12 mg kg ¹, particularly in arid regions with annual rainfall below 550 mm (Hall, 2010). For instance, boron contamination at the Rio Tinto Borax site in California has led to desertification, soil degradation, and loss of vegetation (Kayama, 2010). Excess boron disrupts plant physiology, impairing metabolism, reducing root cell division, lowering chlorophyll content, and ultimately inhibiting growth (Nable et al., 1997; Camacho-Cristobal et al., 2008). As plant biomass declines, organic matter input decreases, further depleting soil fertility and water retention, creating a feedback loop that accelerates land degradation (Parton et al., 1987). Despite the critical role of soil microbial communities in maintaining soil health, the below-ground impact of boron accumulation remains poorly understood. High boron concentrations have been linked to reduced microbial respiration and decreased fungal diversity (Vera et al., 2019). However, the broader effects on bacterial community structure and diversity remain largely unexplored.

In this study, desertification-like conditions were experimentally simulated in boron- contaminated Aquic Ustifluvent (AU) agricultural soil through the application of boron-based fertilizers followed by prolonged exposure to natural environmental conditions, including sunlight, temperature fluctuations, humidity, and rainfall, for a period of 365 days. This long-term exposure was designed to mimic the gradual physicochemical deterioration and environmental stress commonly associated with soil degradation and desertification processes in agricultural ecosystems. To understand the ecological impact of boron toxicity on soil microbial dynamics, a comparative investigation was conducted between boron-contaminated and uncontaminated AU soils. High-throughput culture-independent metagenomic approaches were employed to comprehensively analyse shifts in the bacterial community structure, diversity, and composition under boron-induced stress conditions. The study particularly focused on identifying microbial taxa that were either suppressed or selectively enriched in response to elevated boron concentrations and prolonged environmental exposure. The findings revealed significant alterations in the soil microbial community, indicating that boron contamination exerts strong selective pressure on native bacterial populations. Such microbial shifts may influence critical soil functions, including nutrient cycling, organic matter decomposition, stress adaptation, and overall soil fertility. Furthermore, the study provides important insights into the role of microbial communities in the progression of soil degradation and highlights the adaptive potential of boron-tolerant microorganisms in stressed agricultural soils. These observations contribute to a better understanding of microbe–soil interactions under boron stress and may support the development of sustainable strategies for soil health restoration and management in boron-affected regions.

## 2. Materials and methods

### 2.1. Selection of suitable soil sample to mimic boron-induced desertification for studying soil-bacterial- communities compared and contrasted with the unamended counterpart

Two important criteria were chosen for this study: (i) soil from agricultural field where boron is amended (to cultivate vegetable crops, like cauliflower, that have high boron demand), supposedly would enable isolation of boron tolerant bacteria; and (ii) essentially the soil sample should correspond to semi-arid type of soil having low water storage capacity and relatively high water transmission characteristics. Samples, MA1 (26°31′50.7″N, 88°50′09.8″E) and MA2 (26°31′51.5″N, 88°50′13.5″E) were collected from one location in Maynaguri, within the Jalpaiguri district, West Bengal, India (Figure 1). Soil samples were collected from boron amended and unamended agricultural fields using sterilized containers, gloves, spatula and were transported to the laboratory immediately after sampling to start the downstream experiments within 24-48 h. The boron content of the soil samples was determined by extraction of boron from soil by 2-ethyl-1,3-hexanediol based extraction protocol followed by curcumin based spectrophotometric method ( Yamada and Hattori, 1986; Wimmer and Goldbach, 1999). The soil-types of each sampling sites were decoded from the soil map where the mapping was done in 1:250,000 scale and published in 1:500,000 scale covering the entire state of West Bengal, India (Figure S1 and S2) (Ref: Soils of West Bengal for optimizing land use. 1992. NBSS Publ. 27. National bureau of soil survey & land-use planning *in coop* with Department of Agriculture, West Bengal and Bidhan Chandra Krishi Viswa Vidyalaya, West Bengal. ISBN: 81-85460-12-4.).

**Figure 1.**
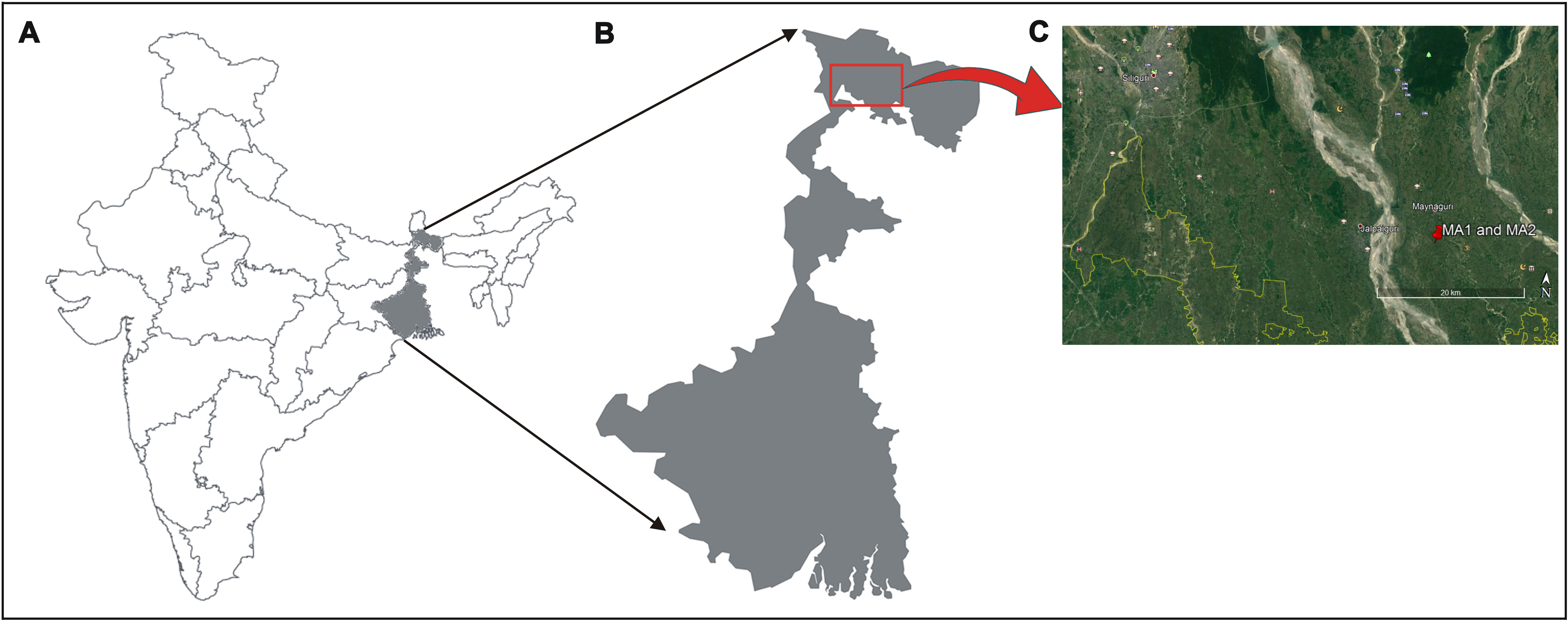
Geographical location of the sampling sites: **(A)** Map of India, showing the overview of the sampling state (**B**) Map of West Bengal, showing the overview of the sampling sites. (**C**) Satellite view of the sampling sites MA1 and MA2 with exact geographical coordinate location.

The soil samples, MA1 and MA2 were classified as aquic ustifluvent—representing a semi-arid soil type (transitional between udic and aridic conditions) and aquic udifluvaquents, which corresponds to a humid soil type that remains moist in most years. MA1 and MA2, having a shallower water table, and material in the upper 15 cm being rested on fine stratified sediment, used as cropland which is irrigated, was chosen to mimic boron-induced desertification. The soil samples (depth ∼15 to 20 cm) were collected following a spatially stratified, and random sampling approach. Each of the two plots was divided into equal sectors of 1 square meter. For each plot, equal amount of fresh soil samples was collected randomly from 3 different points. Immediately after sampling, soil samples were transported to the laboratory. Three soil samples per plot were evenly mixed. Two different pots were filled with equal amounts of soil samples from MA1 (boron-unamended) and MA2 (boron-amended) respectively. Soil-filled pots were placed in open space, (26°42’38.5"N, 88°21’06.4"E), exposed to natural environmental conditions (temperature, sunlight, humidity, and rainfall; Table S1) and observed for next 365 days for primary plant succession in MA1 and MA2 soil-filled pots; and on 0th day, 150th day and 365th day soil - metagenomic DNA was isolated to study bacterial community differences and diversities.

### 2.2. Metagenome extraction and analysis of bacterial diversity

Total community DNA was extracted from soil samples using the PowerSoil DNA Isolation Kit (MoBio, USA). The V3 region of bacterial 16S rRNA genes were PCR amplified using domain-specific universal oligonucleotides from the extracted metagenome, following the fusion primer protocol (Roy *et al*., 2020; Mondal *et al*., 2022); and the PCR products were sequenced on Ion S5 system (Thermo Fisher Scientific, USA). The sequenced files were deposited to Sequence Read Archive of National Center for Biotechnology Information, under BioProject accession number <u>PRJNA642224</u>, <u>PRJNA642225</u> and <u>PRJNA733696</u>, with distinct run accession numbers (Table S2). The low-quality reads, polyclonal reads, barcode sequences and adaptor sequences were trimmed by inbuilt PGM software of Ion S5 system. The filtered sequence reads were again filtered for quality value 20 and sequence length threshold 100 bp. Then, they were clustered into operational taxonomic units (OTUs) at the 97% 16S rRNA gene sequence similarity level, using a range of modules of UPARSE (Edgar, 2013); singletons were excluded at the time of final OTU preparation (Table S2). Determination of Shannon and Simpson indices were done manually. Taxonomic affiliation of the consensus sequence of each OTU was determined using RDP Classifier (http://rdp.cme.msu.edu/classifier/classifier.jsp) at a confidence level of 80%. Rarefaction analysis (using MG-RAST) for all the sequence datasets confirmed that their read-contents were sufficient to reveal most of the diversities present in the samples (Figure S3).

### 2.3. Metagenomic data analysis by MicrobiomeAnalyst

Further analysis was performed in MicrobiomeAnalyst where data was deposited in form of OTU abundance table for marker data profiling (Chong *et al*., 2020). 20% sample prevalence was applied to filter the low count features and 10% inter-quantile range was applied to filter low variance data to remove sequencing errors or low level contaminations. Data was rarefied to the minimum library size and total sum scaling was applied to scale the data. Beta diversity was measured using Bray-Curtis distance method and PERMANOVA was used as statistical method to construct the PCoA plot. Hierarchical Clustering & Heatmap Visualization was performed at genus level using Euclidean distance measure and ward clustering algorithm. Dendogram analysis was carried out using Bray-Curtis index at feature level. Differential abundance analysis was executed using edgeR algorithm and *p-value* cut off was adjusted to 0.05 (Dhariwal *et al*., 2017).

## 3. Results

### 3.1. Soil sample characteristics relative to texture, pH, temperature and boron contents

MA1 and MA2 soil samples were collected from Jalpaiguri district of West Bengal, India. After the collection on different day (0th, 150th and 365th) soil texture, pH, temperature and boron concentration were determined and it was found that the texture of MA1 and MA2 were loamy type. The MA1 and MA2 soil samples were analysed on different days (0th, 150th, and 365th) for soil texture, pH, temperature, and boron concentration, and it was found that both samples had a loamy texture. All the samples were slightly acidic in nature (pH ≤6.5) and the temperature of soil samples was between 21 and 28 °C (Table 1). Among two soil samples, MA2 contain higher amount of boron (90±15.0 ppm in 0 day, 75±12.0 ppm in 150 day and 08±1.54 ppm in 365 day) and MA1 contain less boron (only 03±0.5 ppm in 0 day, 02±0.46 ppm in 150 day and 02±0.53 ppm in 365 day) which was in the normal range of boron concentration in soil (Table 1).

**Table 1.**
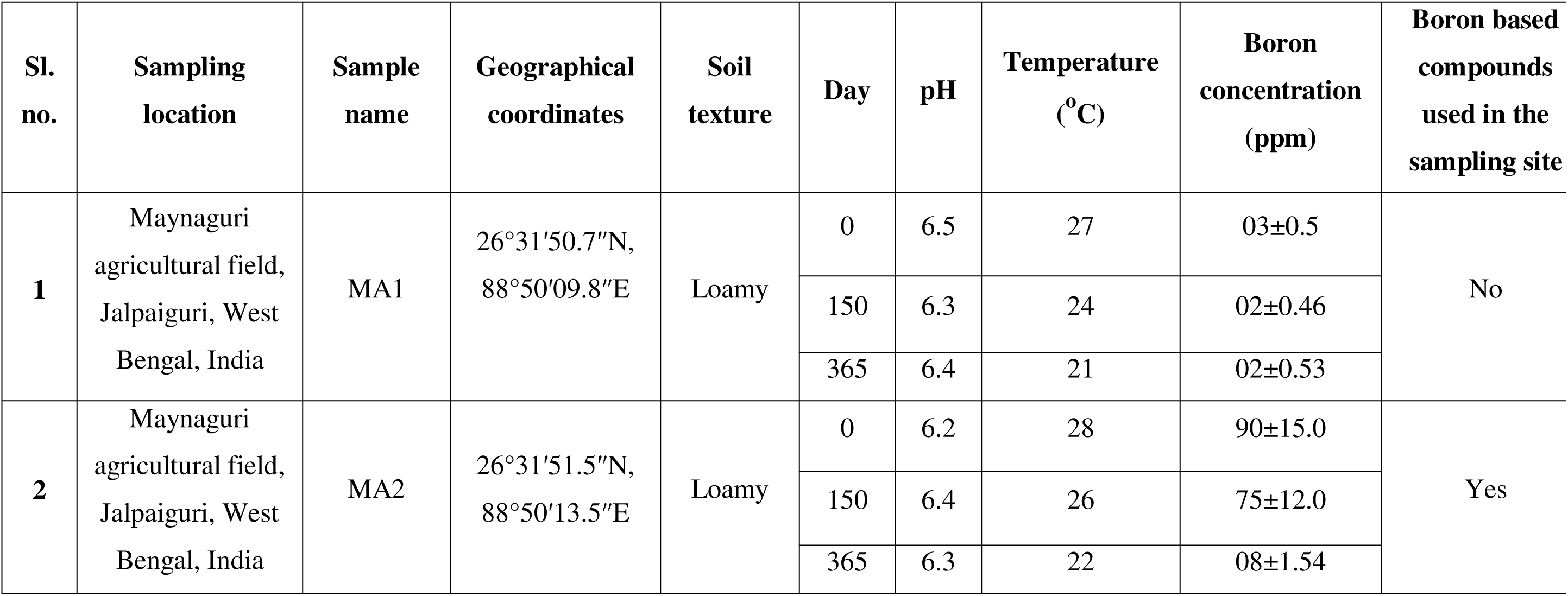
Details of the soil sampling sites along with physicochemical properties.

### 3.2. Overground demonstration of barrenness of boron-amended soil contrasted with plant-succession in unamended soil

Two different pots, MA1 (boron-unamended) and MA2 (boron-amended) were filled with equal amounts of soil samples (collected from Maynaguri on 8 January 2019) and placed in open space for 365 days, from 09 January 2019 to 08 January 2020. Both sites experienced seasonal variation in rainfall (∼1 - 804 mm), temperature (∼16.7 - 27°C), and humidity (∼36 - 92%), peaking during June - September (Fig. 2A). The weather data has been authenticated from https://www.worldweatheronline.com/bagdogra-weather/west-bengal/in.aspx (Table S1). On the 0th day, no seedling was seen in MA1 and MA2 pots. On the 15th day, grass seedlings, constituting the majority, were too immature to identify, along with a few identifiable seedlings of *Cyperaceae*, were observed in the MA1 pot; while MA2 was devoid of any such seedlings. Observations in the MA1 pot on 30th and 45th days were similar to those of the 15th day observation, except there was an increase in length of the young plants; the MA2 pot remained non-green. Since the transition from 45th day to 60th was accompanied by a decrease in humidity in the month of March, the plants growing in MA1 pot had shown signs of wilting due to water-stress. Some plants belonging to *Poaceae* and *Cyperaceae,* and one plant belonging to *Commelinaceae* were visible. In addition to all monocotyledonous plants, only one seedling of a dicotyledonous plant was visible. On the other hand, MA2 pot did not show any sign of plant growth whatsoever on the 60th day of observation. Taken together, the observations on 75th and 90th day(s), it was found that the population of broad-leaved grasses has reduced (may have died due to water stress), while members of the *Cyperaceae* dominated; one plant, *Cyanotis* sp. (family: *Commelinaceae*) and six plantlets of *Pouzolzia indica* (family: *Urticaceae*) thrived on MA1 pot. No signs of plant growth were observed in MA2 pot. On 105th day of observation, MA1 pot has shown one clearly identifiable *Pouzolzia indica* with 3-4 branches, and dominated population of *Cyanotis*/*Murdannia*, while pot MA2 were still without any plants. On 120th day, *Pouzolzia indica* was shown to develop flowers, and plants belonging to Cyperaceae were dominating, while many *Cyanotis*/*Murdannia* (family: *Commelinaceae*) were also seen. Strikingly, the first sign of plant growth, comprising of three very young seedlings, belonging to *Cyperaceae* family, appeared in MA2 pot on the 120th day, and grew to eight visibly identifiable individual *Cyperus* sp. on 150th day of the experiment. On 150th day, MA1 pot had demonstrated profuse growth of *Pouzolzia indica*, Cyperus could be identified as Cyperus compressus, and *Cyanotis*/*Murdannia* plants had shown branching, but they were yet to flower. On the 300th and 365th day both pots contain mostly similar types of plants with some exceptions (Fig. 2B & 2C).

**Figure 2.**
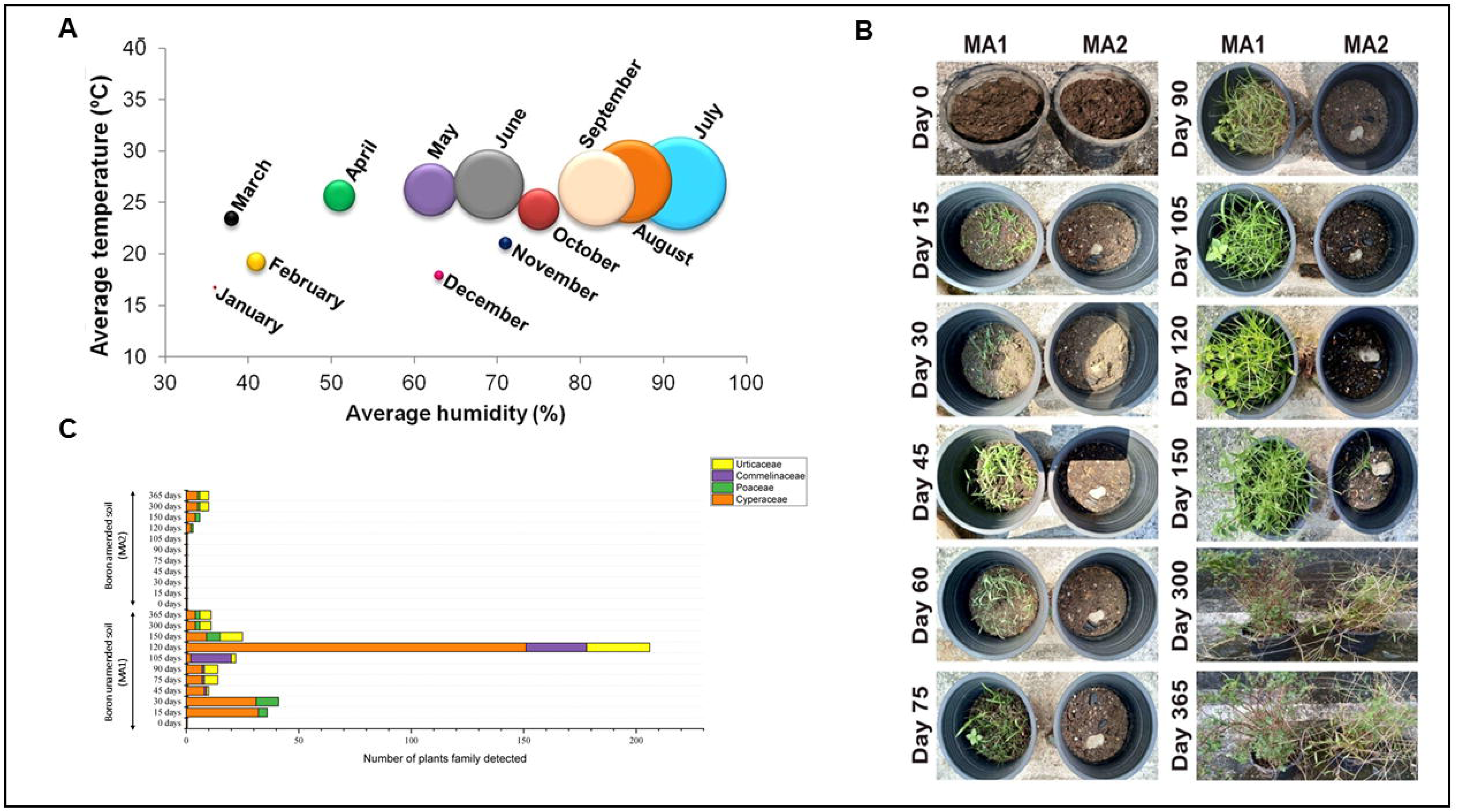
**(A)** Bubble plot comparing the temperature, humidity and rain fall of the different months when the experiment was carried out. Humidity is plotted along the X-axis, temperature along the Y- axis, while individual months are represented by different colour circles. Size of the circles is proportional to the amount of rainfall. **(B)** Type of natural flora was observed in MA1 (boron unamended soil) and MA2 (boron amended soil) sample on different day throughout the experiment. **(C)** Graphical representation of type of plant species found in MA1 and MA2 samples throughout the experiment.

### 3.3. Microbiome analysis of boron amended and unamended soil

Frequent and excessive use of boron fertilisers in agriculture can contribute to desertification when it affects the natural succession of plants, causing a decline in soil microbial diversity. We hypothesized that under selective pressure, resulting from soil amendment with boron, and withdrawn replenishment of soil nutrient from decomposed plant parts of the overground vegetation, only a sub-set of soil bacteria (that thrived in the same unamended soil) comprising of bacteria capable of degrading a broad spectrum of carbon and energy sources and electron acceptors, members of the phyla that can form spore under nutrient-scarce conditions, bacteria capable of tolerating high boron, low-nutrient condition (oligotrophic condition), and soil-inhabiting photoautotrophs would sustain and survive. To test our hypothesis, on 0th, 150th and 365th day, soil-metagenomic DNA was isolated to study bacterial community differences and diversities. MA2 sample had higher boron concentration in comparison to the unamended MA1 soil sample (Table 1). Bacterial metataxonomic analyses (based on V3 region of 16S rRNA gene) revealed a contrasting picture of bacterial diversity (boron unamended vs amended soil) (Table S2). The alpha diversity indices were calculated to evaluate the microbial diversity of boron amendend (MA2) and unamended (MA1) soil samples. According to the Simpson reciprocal index, in case of boron amended condition 365th day soil sample had the most diversity, followed by 150th and 0th day. On the other hand in case unamended soil 0th day soil sample had the most diversity followed by 365th and 150th day. According to Shannon’s diversity index (H), in case of boron amended condition the highest level of community heterogeneity was found in 365th day soil sample, followed by 150th and 0th day. On the other hand in case of unamended soil the highest level of community heterogeneity was found in 365th day soil sample, followed by 0th and 150th day. Evenness was maximum in 365th day soil sample, followed by 150th and 0th day according to Shannon’s equitability index (ESI), in case of boron ameded condition. On the other hand in case of unamended soil evenness was maximum in 365th day soil sample, followed by 0th and 150th day according to Shannon’s equitability index (ESI) (Table S2). In comparison, beta diversity analysis revealed that the boron amended and unamended soil samples were obviously dissimilar in terms of microbial composition, whereas the 150th and 365th day soil samples were less dissimilar compare to the soil samples of 0th day (Fig. 3A). In case of MA1 sample at 0th to 150th day ≈22% OTUs increment was observed; on the other hand in MA2 sample in the same time frame only ≈16% OTUs increment was observed. Interestingly at 150th to 365th day ≈27% OTUs increment was observed in MA2 sample, but in MA1 sample ≈35% OTUs was decreased. Maximum difference between OTUs was observed in 150th day. The number of OTUs detected in MA1 (3190 OTUs) was higher than MA2 (2022 OTUs), indicating ≈37% decrease in OTUs in soil amended with boron in 150th day. In all the boron amended samples, a total of 6504 OTUs were detected, which were distributaed in 21 different phyla. The majority of sequences belonged to members of *Firmicutes* (26%), *Proteobacteria* (18%), *Actinobacteria* (14%) and *Acidobacteria* (6%) (Fig. 3B and Table S3). In boron amended soil the phyla *Firmicutes* (in 0th day- ≈42%, 150th day- ≈27% and 365th day- ≈16%), *Proteobacteria* (in 0th day- ≈8%, 150th day- ≈20% and 365th day- ≈22%), *Actinobacteria* (in 0th day- ≈19%, 150th day- ≈11% and 365th day- ≈13%), *Acidobacteria* (in 0th day- ≈3%, 150th day- ≈5% and 365th day- ≈8%) and *Bacteroidetes* (in 0th day- ≈0.4%, 150th day- ≈3% and 365th day- ≈5%) are the most abundant (Fig. 3B). In all boron unamended samples, a total of 8037 OTUs were detected belongs to 21 different phyla. The majority of sequences belonged to members of *Firmicutes* (15%), *Proteobacteria* (23%), *Actinobacteria* (14%) and *Acidobacteria* (7%) (Fig. 3B and Table S3). In boron unamended soil the phyla *Firmicutes* (in 0th day- ≈17%, 150th day- ≈18% and 365th day- ≈10%), *Proteobacteria* (in 0th day- ≈23%, 150th day- ≈22% and 365th day- ≈23%), *Actinobacteria* (in 0th day- ≈19%, 150th day- ≈10% and 365th day- ≈14%), *Acidobacteria* (in 0th day- ≈5%, 150th day- ≈8% and 365th day- ≈9%) and *Bacteroidetes* (in 0th day- ≈6%, 150th day- ≈5% and 365th day- ≈6%) are the most abundant (Fig. 3B). The phylum *Armatimonadetes, Balneolaeota* and *Fusobacteria* only observed in boron amended condition. In boron amended condition (MA2) a increment in OTUs was observed for the phylum *Acidobacteria*, *Bacteroidetes*, *Candidatus Saccharibacteria*, *Chloroflexi*, *Gemmatimonadetes*, *Nitrospirae* and *Proteobacteria*. In the MA2 sample a initial decrease in OTUs observed for the phylum *Actinobacteria* and *Cyanobacteria*, but the OTUs again increase in 365th day. In case of *Firmicutes* OTUs deduction rate was very slow and comperatively higher number of OTUs was observed in 0th, 150th, 365th day of boron amended soil (Fig. 3B and Table S3).

**Figure 3.**
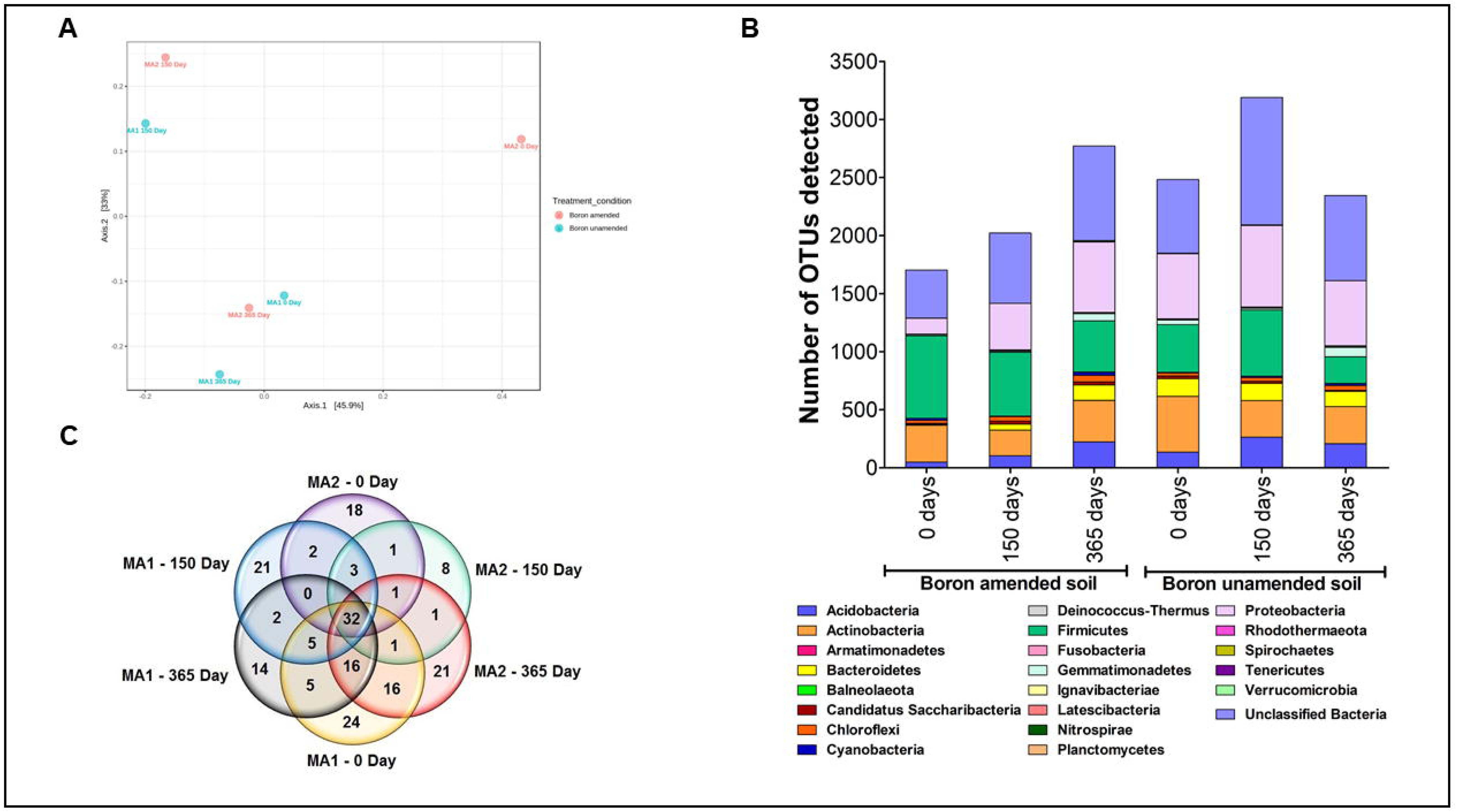
**(A)** Principal coordinate analysis plot of all samples between boron amended (represented as MA2) and unamended (represented as MA1) soil. Each dot represents one sample of different day throughout the experiment. **(B)** Microbial community in MA1 (boron uamended soil) and MA2 (boron amended soil) on different days of experiment; phylum level distribution of the OTUs was plotted here. **(C)** Venn diagram displaying the common and unique genera among all six (boron amended and unamended) soil sample microbial communities.

A increment in bacterial genera was detected in MA2 metagenome (0th day - 132, 150th day - 145 and 365th day - 218); on the other hand, in MA1 sample total number of bacterial genera are more or less similar in different days (0th day - 202, 150th day - 187 and 365th day - 184). Boron rich desertified soil sample, MA2, contained 47 exclusive genera (18 on 0th day, 8 on 150th day and 21 on 365th day sample) (that were not present in MA1), while 59 genera were exclusively present in MA1 soil sample (24 on 0th day, 21 on 150th day and 14 on 365th day sample) (absent in MA2 soil sample) (Fig. 3C). From the relationship study it was also found that 32 genera were found in all the soil samples (Fig. 3C). Among the exclusive taxa, besides several organotrophic genera, *Kocuria,* reported to grow under iron limitation; *Armatimonadetes,* defined as aerobic soil-based oligotroph; *Laceyella,* reported as non-acid fast endospore former; and *Cetobacterium* and *Oribacterium,* reported as anaerobic bacteria, were present in the metagenome of MA2. While segregating OTUs as per affiliation to Gram-positive and Gram-negative taxa (excluding the Gram-variable ones), it was found that MA2 exceeded MA1 by ≈4% (≈34% in MA1; ≈38% in MA2) in terms of percentage distribution of Gram-positive taxa, and MA1 exceeded MA2 (≈65% in MA1; ≈62% in MA2) by ≈3% in terms of percentage distribution of Gram-negative taxa (Table S4). The information yielding from the extensive survey of literature on culturable boron-tolerant bacteria, when superimposed on the data obtained from this study, 12 bacterial taxa, *Acinetobacter*, *Alkaliphilus*, *Anoxybacillus, Arthrobacter*, *Bacillus*, *Gracillibacillus*, *Lysinibacillus*, *Microbacterium*, *Oceanobacillus*, *Pseudomonas*, *Ralstonia* and *Streptomyces,* having report to tolerate varying concentrations (10 mM to- 450 mM) of boron have been identified, and nine of them were Gram-positive (Table S4) (Ahmed *et al*., 2007a-d; Miwa *et al*., 2008; Miwa *et al*., 2009; Yoon *et al*., 2010; Raja and Omine, 2012; Nural *et al*., 2018). We recalled our hypothesis at this point that under selective pressure, resulting from soil amendment with boron, there must be preponderances of oligotrophic Gram-positive boron tolerant bacteria in boron amended agricultural soils of the northern West Bengal, India.

According to the relative abundance analysis we have found that *Brevibacillus, Desulfotomaculum, Gracilibacillus, Gracilibacter, Hyphomicrobium, Laceyella, Ornithinimicrobium, Paenibacillus, Planifilum, Romboutsia* and *Vulgatibacter* are the most abundant genera in the boron amended soil throughout the time frame (0th to 365th day) (Figure S4). Differential abundance at different levels of all the samples were measured by edgeR algorithm in MicrobiomeAnalyst. The result shows that significant difference was found only, for one class Acidobacteria_Gp17 (*p-value*= 0.04533); for one order Geodermatophilales (*p-value*= 0.0049368); for two family Geodermatophilaceae (*p- value*= 0.0049368), Ornithinimicrobiaceae (*p-value*= 0.025242); and for 5 genus *Actinocorallia* (*p- value*= 0.0049368), *Chitinophaga* (*p-value*= 0.049296), *Geodermatophilus* (*p-value*= 0.0049368), *Gracilibacillus* (*p-value*= 0.043642), *Ornithinimicrobium* (*p-value*= 0.025242) (Figure S5).

## 4. Discussion

Boron, an essential micronutrient for plants, is absorbed primarily as boric acid through the root system. However, its optimal concentration range in soil is exceedingly narrow, with toxicity occurring at only a few ppm and deficiency below 0.5 ppm, making boron management in agriculture particularly challenging (Nable et al., 1997; Çöl and Çöl, 2003; Pahl et al., 2005; Gentz and Grace, 2006; Cervilla et al., 2007). Agricultural soils that lack sufficient boron for one crop may exert toxic effects on another, complicating fertilization strategies (Brdar-Jokanović, 2020). Frequent and excessive boron fertilisation disrupts plant succession and soil microbial diversity, contributing to desertification. Despite the well- documented effects of boron toxicity on plant physiology, its impact on below-ground microbial communities remains underexplored.

In this study, we investigated the influence of boron amendment on soil microbial dynamics and primary plant succession in *Aquic Ustifluvent* (AU) agricultural soil. Soil from boron-unamended (MA1) and boron-amended (MA2) conditions was exposed to natural environmental conditions for 365 days, monitoring plant establishment and microbial shifts over time. Initial analyses confirmed significantly higher boron levels in MA2 compared to MA1 (Table 1). Over the first 45 days, MA1 supported early plant colonization, whereas MA2 remained barren. By the 120th day, the first signs of plant growth, primarily *Cyperus* sp., emerged in MA2, contrasting with the diverse plant community established in MA1. By the 365th day, both pots contained plant species from families *Commelinaceae* and *Urticaceae*, though the delay in succession in MA2 suggested a strong initial inhibitory effect of boron on plant establishment. To explore the microbial response to boron stress, metagenomic DNA was extracted from soil at 0, 150, and 365 days. We hypothesized that boron amendment exerts selective pressure on soil microbial communities, favoring taxa capable of surviving high-boron, low-nutrient conditions. Comparative metagenomic analysis revealed a substantial reduction in bacterial diversity in MA2. The proportion of unclassified OTUs, often indicative of taxa with unknown ecological functions, was disproportionately higher in MA1 (classified:unclassified OTUs = 1848:635, 2091:1098, 1613:732) compared to MA2 (1289:414, 1417:605, 1957:815) across time points (Table S2). While unclassified sequences are commonly observed in culture-independent studies, they remain underexplored despite their potential significance in microbial community dynamics.

Despite a reduction or increment in phylum-level OTU, a high diversity of *Acidobacteria* subgroups was observed in boron amended soil sample, MA2, and the highest number of OTU was found for the *Acidobacteria* subgroup Gp6 (Table S4), corroborating the observation by Kielak *et al*., 2016 that *Acidobacteria* subgroup Gp6 responds positively to the high content of boron in soil. A recent study has shown that both *Candidatus Saccharibacteria* and *Chloroflexi*, at the phylum level, were prominent among other specialists that had a significantly higher abundance under varied field sites and fertigation regimes (Xu *et al*., 2020). *Candidatus Saccharibacteria* is known to have a part in the degradation of diverse organic compounds in addition to sugar compounds under aerobic-nitrate reducing-and anaerobic conditions (Kindaichi *et al*., 2016). Since *Chloroflexi* can rely on photosynthesis to produce energy, they can survive in soils of relatively poor fertility (Fullerton and Moyer, 2016), may have added to the increase in OTU reads from boron amended soil. This is indicative of the fact that the physico-chemical status, particularly the bioavailability of carbon and nitrogen, may indicate the predominant strategy of microbial life in the given soil conditions. It is rather probable that due to the natural succession of plants, input of labile carbon adding litter and root exudates, with the latter known to pick up and shape the microbial makeup of the rhizosphere (Leff *et al*., 2015; Ridl *et al*., 2016) may leave an impression on microbial community by changing the composition. Earlier authors have predicted that *Acidobacteria* congregates the supposition of an oligotroph as it prevails in carbon-poor soils as well as in bulk soil where resources, if bioavailable, are present at comparatively lower concentrations than in the rhizosphere (de Castro *et al*., 2013). While *Actinobacteria*, in general, among soil microbial phyla, has been considered to be more copiotrophic, some members of *Actinomyceteles* are capable of depolymerising complex carbon substrates such as lignin or cellulose (Aislabie *et al*., 2013). Therefore, the relatively high abundance of *Actinobacteria* OTUs in MA2 could be due to enhanced breakdown of more refractory carbon substrates, rather than labile ones. There was no OTU representation in the phyla, *Spirochaetes* and *Tenericutes* on 150th day (100% reduction in OTU number), but surprisingly there has been less reduction in OTUs representing *Firmicutes* phylum (members of the phylum show an explicit property to develop heat-resistant endospores under nutrient- deficient conditions) in MA2 compared to MA1. Also, several isolates belonging to genera like *Bacillus, Enterococcus and Lysinibacillus* under *Firmicutes* were reported to tolerate boron (Sen *et al*., 2020). Hence, it is likely that boron tolerant bacteria belonging to the phylum *Firmicutes* might have contributed to their OTU abundance in the MA2 soil having excess boron (Table S4).

Our findings also revealed a shift in microbial community composition, with certain genera disappearing while new taxa emerged. MA2 soil harbored exclusive OTUs from 47 genera, including heavy metal- and metalloid-resistant taxa such as *Acidiferrimicrobium*, *Achromobacter*, *Bhargavaea*, *Bradyrhizobium*, *Frateuria*, *Acidobacteria*, *Idiomarina*, *Luteibacter*, *Mucilaginibacter*, *Kocuria*, and *Nocardiopsis*. These genera have been previously associated with resistance to various metals, including Cd², Zn², Pb², As(V), Cu², Ni², and Cr. Additionally, several genera known for boron tolerance were detected in MA2, including *Acinetobacter* (50 mM), *Alkaliphilus* (10 mM), *Anoxybacillus* (70 mM), *Arthrobacter* (80 mM), *Bacillus* (450 mM), *Gracillibacillus* (50–450 mM), *Lysinibacillus* (50–230 mM), *Microbacterium* (200 mM), *Oceanobacillus* (450 mM), *Pseudomonas* (100 mM), *Ralstonia* (100 mM), and *Streptomyces* (80 mM) (Table S4). The enrichment of boron-tolerant and oligotrophic bacteria in MA2 suggests that boron stress selects for microbial taxa capable of survival under extreme conditions. These adapted communities likely played a role in the eventual establishment of plants in MA2 by contributing to nutrient cycling and organic matter decomposition. The delayed yet eventual colonization of angiosperms in MA2 after three months of barrenness underscores the resilience of soil microbiota in mitigating boron-induced stress. Recognizing the limitations of culture-independent approaches in capturing the functional potential of microbial communities, we complemented our analysis with culture-based methods to assess the physiological traits of boron-tolerant isolates. The integration of both approaches provides a more comprehensive understanding of microbial adaptations to boron stress and their potential role in soil ecosystem recovery. Overall, our findings highlight the profound impact of boron contamination on soil microbial diversity and succession. The selection pressure imposed by high boron levels leads to a distinct microbial community shift, favoring taxa with metal resistance and stress tolerance. Understanding these microbial adaptations is crucial for developing strategies to mitigate boron-induced desertification and restore soil health in affected agricultural landscapes.

## CRediT authorship contribution statement

**Subhajit Sen:** Writing – original draft, Visualization, Methodology, Investigation, Formal analysis, Conceptualization**. Nibendu Mondal:** Writing – review & editing, Visualization, Formal analysis. **Chandana Basak:** Writing – review & editing, Visualization, Formal analysis**. Wriddhiman Ghosh:** Writing – review & editing, Conceptualization. **Ranadhir Chakraborty:** Writing – review & editing, Visualization, Conceptualization, Supervision.

## Acknowledgments

Authors sincerely thank Department of Science and Technology (DST), Government of India as one of the author; Subhajit Sen received research grants through INSPIRE Fellowship vide sanction order DST/INSPIRE Fellowship/2016/IF160786. We also thank Science and Engineering Research Board (SERB), Bose Institute and Council of Scientific and Industrial Research (CSIR), India provided research grants to Nibendu Mondal and Chandana Basak respectively. The support for consumables and experiments was derived from University of North Bengal, Bose Institute and DST contingency.

## Appendix A.Supplementary data

Supplementary data to this article can be found online at

## Data Availability Statement

Soil sample metataxonomy raw read sequence datasets were deposited in the National Center for Biotechnology Information (NCBI) Sequence Read Archive (SRA) under BioProject accession number PRJNA642224, PRJNA642225 and PRJNA733696.

---

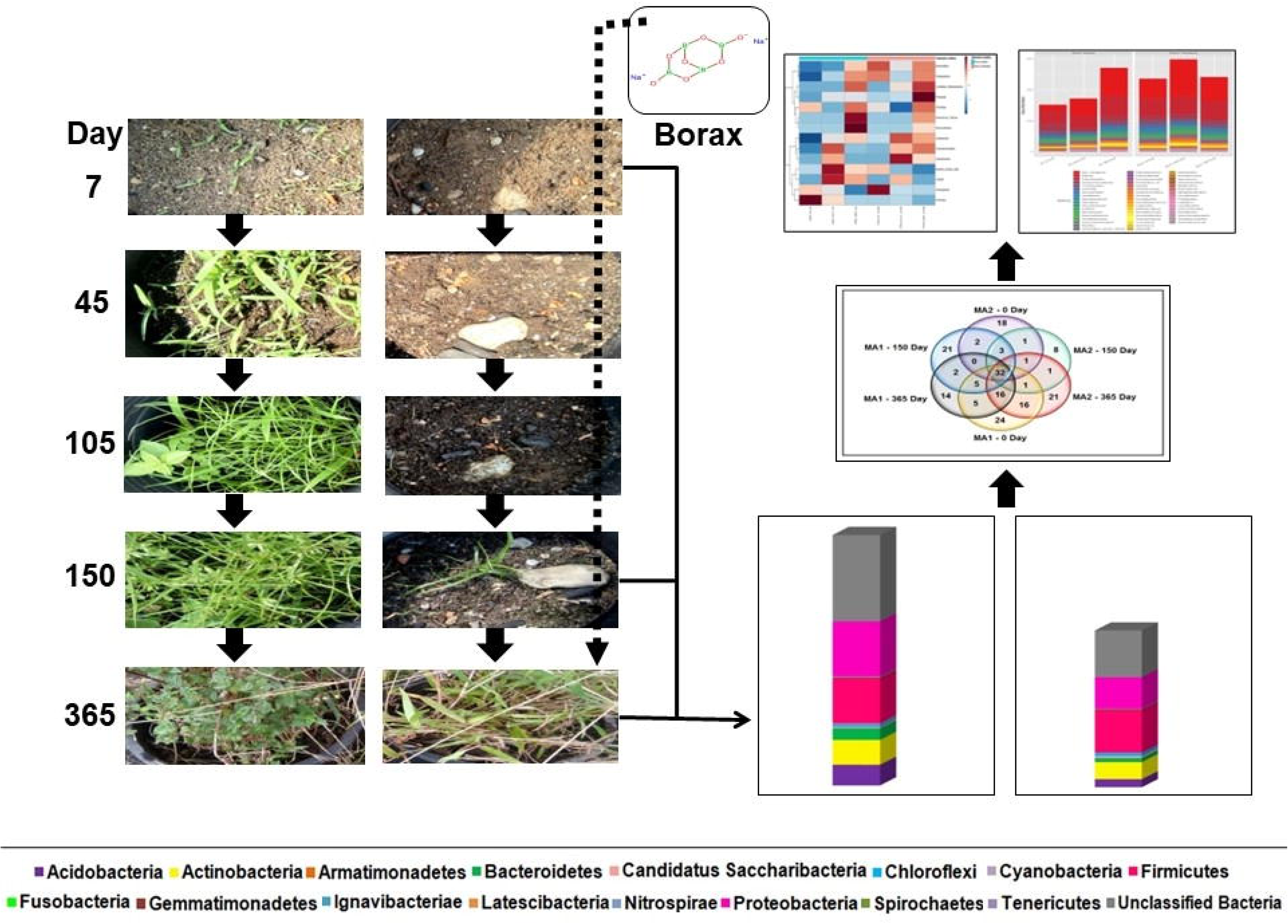

